# Signatures of personality on dense 3D facial images

**DOI:** 10.1101/055590

**Authors:** Sile Hu, Jieyi Xiong, Pengcheng Fu, Lu Qiao, Jingze Tan, Li Jin, Kun Tang

## Abstract

It has long been speculated that there exist cues on human face that allow observers to make reliable judgments of others’personality traits. However, direct evidences of association between facial shapes and personality are missing. This study assessed the personality attributes for 834 Han Chinese volunteers (405 males and 429 females) utilizing the five-factor personality model (the ‘Big Five’ model), and collected their neutral 3D facial images. Dense anatomical correspondence was established across the 3D facial images to allow high-dimensional quantitative analyses on the face phenotypes. Two different approaches, Composite Partial Least Square Component(CPLSC) and principle component analysis (PCA) were used to test the associations between the self-tested personality scores and the dense 3D face image data. Among the five personality factors, Agreeableness and Conscientiousness in male, and Extraversion in female were significantly associated to specific facial patterns. The personality-related facial patterns were extracted and their effects were extrapolated on simulated 3Dfacial models.

## Introduction

As one of the most complex anthropological traits, human facial shape is strongly regulated by many factors such as genetic inheritance, ethnicity, age, gender and health, etc. Own to the rapid progresses of face imaging and analysis technologies, especially the 3D dense face model based approaches, complex facial shape traits were continuously discovered to signal genetic polymorphisms [1], ethnicity [2], gender [3], diseases [4], health[5], as well as aging[6].

Apart from anthropological perspectives, human face was also intensively studied for their social attributes, such as attractiveness [7–8] and personality [9–10]. The facial signals of social attributes are more difficult to study due to the involute and highly subjective nature of the social attributes. Many people believe that there exist facial cues towards the hidden personality of unknown individuals [11]. Indeed, a series of studies have been carried out to formally test the hypothesis of facial signals of personality. Most of these studies utilized the five-factor model of personality, or the‘Big Five’(BF) model for short [12–14]. The BF model ascribes the personality into five dimensions: extraversion (E), agreeableness (A), conscientiousness (C), neuroticism (N) and openness (O). According to the definitions, a higher score of E, A, C, N and O indicates the character of being outgoing or energetic, friendly or compassionate, self-disciplines or organized, sensitive and nervous, and inventive or curious respectively. The BF model is well suited for the purpose of evaluating the facial signals of personality, due to several advantages. First, the five personality factors were found to be approximately orthogonal to each other [15], as is a desirable statistical property in factor analyses. Second, people’s BF test scores are highly stable during adulthood [16]. In fact, genetic studies revealed relatively high heritability (42∼57%) of the five personality traits [17], suggesting that the BF traits reflect more constitutional characters rather than transient emotional changes. Finally, the BF model have been successfully applied in different genders [18], a variety of cultures [19–20], and even in chimpanzees [21], showing its strong cross-group robustness and compatibility.

Passini and Norman first conducted a seminal study in 1966, in which a small group of volunteers, un-known to each other, were asked to rate themselves and their peers on the BF scales without verbal communication [22]. They found that for extraversion, agreeableness and openness, the self-reported scores and those given by observers were significantly correlated. Other similar studies also confirmed that people can correctly reckon the personality of unknown people to certain extents, at first encounter without verbal communication [11, 23]. Later studies noted that the cues for personalities could be recovered in mere static face pictures with neutral expression [10, 24–25]. To directly obtain the specific facial patterns associated with personality traits, Little and Perrett proposed an ingenious method [25]. They ranked the head portraits of the volunteers along the five BF dimensions by their self-rating,and synthesized the composite portraits for the extreme scorers. They found that raters could identify the BF traits underlying the composite images better than chance, particularly for conscientiousness and extraversion and agreeableness. Similar studies confirmed the accuracy of composite images in guiding the recognition of personality traits [5, 10].

Nonetheless, all of the previous studies relied on the subjective judgment of human raters to evaluate the effects of particular facial signals. A direct association between facial shape changes and the personality is missing. In other words, we still do not know whether the inner personality does induce substantial modulations on individual’s physical facial appearance; and if yes, what are the exact physical changes pertaining to each personality attribute. This question is particularly important to the feasibility of automatic face recognition systems that may judge personality without human interference. Furthermore, all the previous studies were conducted in samples of relatively small sizes. In the current study, we followed a novel strategy by utilizing 3D images to uncover the association between the human face and personalities. Inbrief, we collected 3D facial images of 834 volunteers from Shanghai, China using highresolution 3D camera system (3dMD Face System, www.3dmd.com), as well as their corresponding BF questionnaires, in which the result can be transferred to the consecutive score for each BF factor. Thereafter we proposed a partial least square (PLS) based statistical method, named composite PLS component (CPLSC), to inspect the association between the human face and each of BF factor based on the 3D image data and BF scores, and extract the personality-related facial features from the high density 3D image data. In addition, we validated the consistence of results from CPLSC by further principle component analysis (PCA). All the extracted personality-related features were finally visualized and animated by our R package“3DFace”.

## Results

### Personality test using BF model

Personality scores were measured for all the volunteers using a self-report questionnaire (Chinese Version) according to the Big Five Inventory. Cronbach’s αcoefficient was used to evaluate the reliability of the BF scores [26]. Cronbach’s α reflects the correlation of different tests towards the same personality factor to be measured. In our survey, the Cornbach’s α scores ranged from 0.68 to 0.80 with the mean of 0.74 ((Table 1), consistent with the previous surveys in East Asians [27].

**Table 1.** The The Cornbach’s α in each of BF factor in two genders

| Personality | Male | Female |
| --- | --- | --- |
| Extraversion | .80 | .79 |
| Agreeableness | .71 | .68 |
| Conscientiousness | .72 | .73 |
| Neuroticism | .70 | .71 |
| Openness | .76 | .77 |

### Association analyses of 3D facial images and personality based on Partial Least Square (PLS) regression

PLS is a statistical approach to find the maximum potential associations between two variables, either or both of which could be multi-dimensional. It is therefore well suited for finding the latent relationships between the BF factors and the high-dimensional facial morphological data. To control for the gender effect, we conducted the analyses in males and females separately throughout the study. Within each gender group,we carried out PLS regressions for each of the five BF factors separately, and a leave-one-out (LOO) procedure was applied to cross-validate the PLS models (see methods). Similar to principle component analysis (PCA), PLS can also iteratively decomposes the high-dimensional data space into consecutive PLS components (PLSC) [28]. An optimal PLS model is the one composed of the top-ranking PLSCs that can best predict independent data. For each BF factor, we analyzed up to top 20 PLSCs, and coefficient of determination (*R*^2^) was calculated on the cross-validation data to evaluate the accuracy of prediction (Figure 1a and 1b, Method). Positive *R*^2^ values indicate effective predictions and greater *R*^2^ values stand for better performance; on the contrary, an *R*^2^ curve constantly below zero indicates lack of association signals and thus no prediction power towards the corresponding personality trait. We define the effective PLSC number as the number of top PLSCs that rendered the maximum positive *R*^2^ value. As can be seenin (Fig 1a and (1b, the effective PLSC numbers are 2, 3, 3, 2 for E, A, C and N in males; and 2 for E in females respectively. For the other traits, *R*^2^ values were always negative and they were dropped from further PLS analyses.

**Figure 1.**
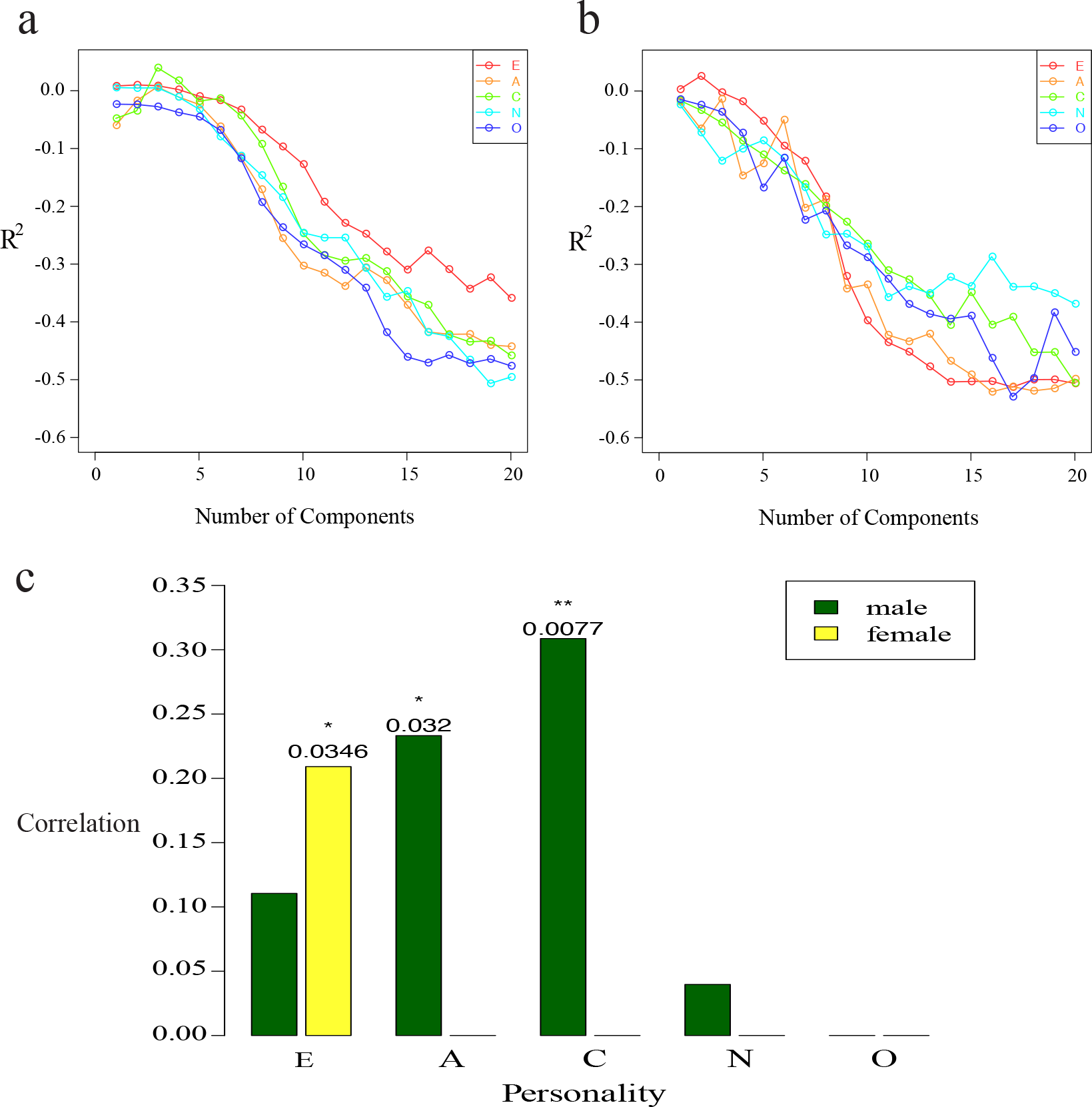
**Determining the effective number of PLSCs for each BF factor, and personality traits significantly correlated between predicted and true scores**. (a) *R*^2^evaluation to determine the effective number of PLSCs number in male samples. (b). *R*^2^ evaluation to determine the effective number of PLSCs in female samples. (c). Personality traits with significant correlation between the predicted and true scores.

For the personality traits that can be effectively predicted, we obtained the Pearson correlation ρ between the true BF scores and the corresponding predicted scores from the LOO procedure, under the optimal prediction models. To formally test the statistical significance, these correlation scores were compared to null distributionsgenerated by re-shuffling the BF scores among different individuals (see Methods for details). As shown in Fig.1c, we found that the 3D facial shapes were significantly associated with personality trait A (Pearson’s correlation ρ=0.2332, *P*value=0.032) and C (Pearson’s correlation ρ=0.3087, *P value*=
0.0077) in males, and E (Pearson’s correlation ρ=0.2091,*P value*=0.0346) in females. For the personality trait E (Pearson’s correlation ρ=0.1105,*P value*=0.147) and N (Pearson’s correlation ρ=0.0397, *P value*=0.3338) in males, although the correlations were positive, the reshuffling test did not support their statistical significance, suggesting that such moderate positive correlations may appear merely by chance.

For the personality traits showing significant correlations with facial shapes, it is desirable to extract the personality related facial signatures, as well as to visualize effects the personality traits induce on face. We designed a method termed the composite PLS components (CPLSC) to extract the personality-related facial signatures from the optimal PLS prediction models (see Methods for details). CPLSC basically identifies a single dimension in the 3D face data space that defines the overall shape changes along with increasing personality score. The effects of such facial signatures may be clearly visualized by extrapolating the mean face to extreme extents in opposite directions along the CPLSC dimension (Fig 2. See Methods for details). The results of A, C in male and E in female are shown in Fig 2. The heat color portraits indicate contribution of each vertex to the shape changes (Methods), with warmer colors signaling greater changes along the CPLSC dimension, and colder colors for minor changes. For agreeableness in male, as shown in the top panel of Fig 2,the CPLSC signature mostly involved an upper facial region around the forehead and eyebrows and a lower region centered on the lower lip. When we gradually change the mean face along the CPLSC dimension towards higher agreeableness (see Fig 2, and Supplementary video), the eyebrows seem to be lifted up, with a reduced forehead span (distance between the eyebrows and the hairline); A more recognizable expression appears around the mouth, where the lips laterally extend outwards and bend upwards at the lip corners, showing a clear expression of“smile”. When the face is morphed towards lower agreeableness, opposite changes happen, including the sinking eyebrows and jaw; the lip corners also drop downwards to give a sign of unhappiness. As for conscientiousness in males, the face of higher C score shows lifted and laterally extended eyebrows, as well as wider opened eyes. Changes in lower facial area mainly involve withdrawing upper lip and pressed muscles around the jaw, showing a pose of“tension” around the mouth area. These are in contrast to the seemly relaxed face associated with low C score, whose brows and eyes were naturally pulled down by gravity, and the muscles around mouth seem to be rather relaxed. It is interesting to notice that the faces of low A and low C scores are similar in the general trend: both faces showed a sense of relaxation and indifference.

The only personality trait found significantly associated in females is extraversion. As shown in Fig 2, The CPLSC model indicates that substantial signals come from around the nose and upper lip. The regions defining the face contour, including the temples, masseters and chin also seem to contribute tothe shape differences. When morphed along the CPLSC dimension, the face of higher E score showed a more protruding nose and lips, and recessive chin and masseter muscles. The face of lower E score showed clearly the opposite changes, whose nasal-maxillary region seems to press against the facial plane.

**Figure 2.**
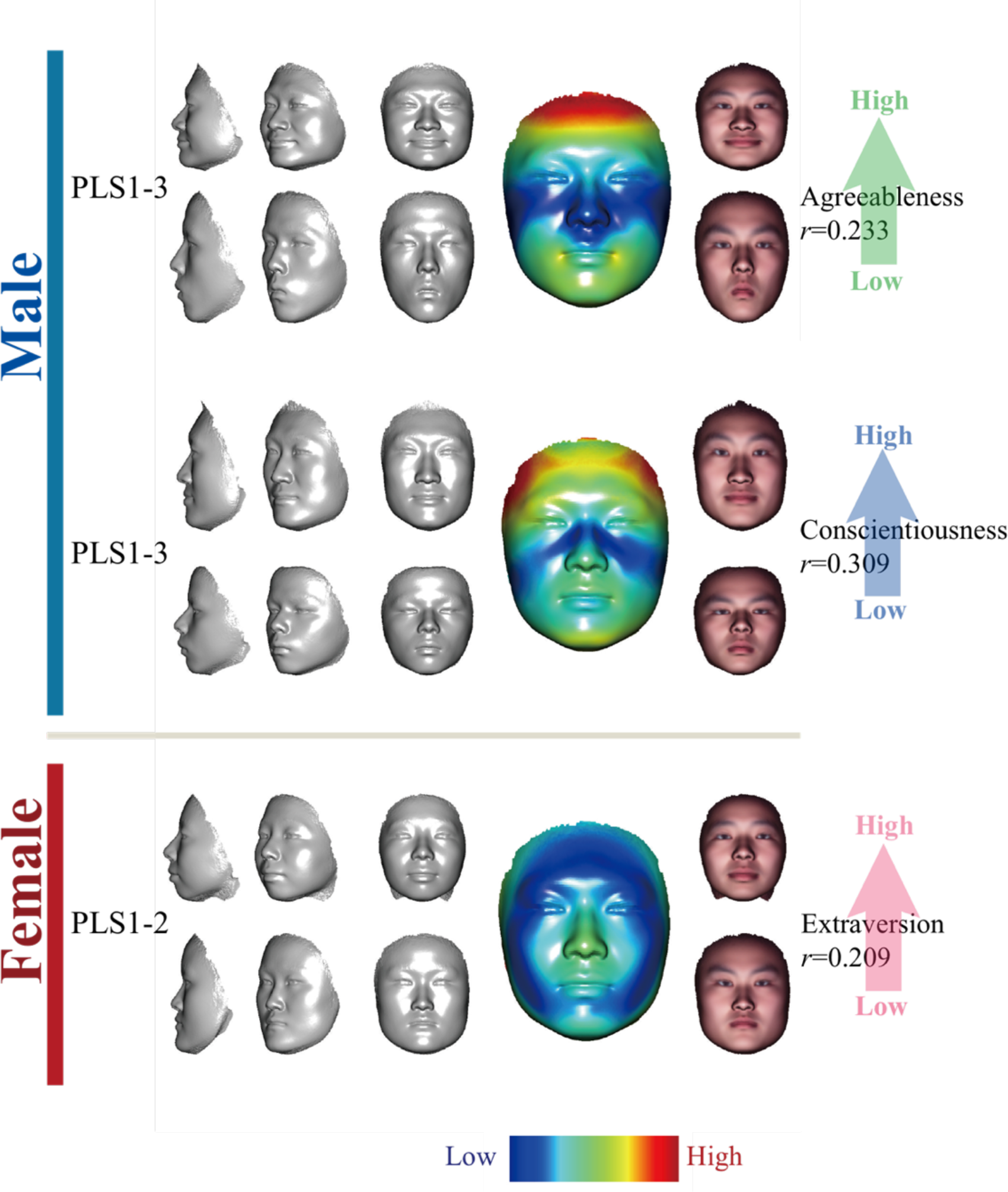
**Features selected by CPLSC model from faces significantly associated with BFfactors intwo gender.** Three panels are ordered from top to bottom sequentially: Agreeableness-related CPLSC which consist of the first three PLSC in male; Conscientiousness-related CPLSC which consist of the first three PLSC in male; Extraversion-related CPLSC which consist of the first two PLSC in female. In each panel, the upper faces are simulated by adding five standard deviation of projected samples to the mean face, the lower faces are created by subtracting standard deviation of projected samples to the mean face. From left to right are faces rotated by 90°, 45° and 0°. The bigger face next is the mean face, on which the heat colors represent the norm value of CPLSC at each vertex. At the right of the mean face, there are two faces the same as the faces at the left of mean face, but with texture.

### Association analyses of 3D facial images and personality based on PCA

PCA provides another way of testing the potential associations between high-dimensional face data to the personality traits. The face data was first decomposed by PCA. Only the top 20 PCs were used in the subsequent analyses, which composed the majority (96.7% in males and 96.1% in females) of the shape variance. Linear regression was used to find the prediction model of personality based on all 20 face PCs, and a similar LOO procedure was used to cross-validate the association (see methods). Applying the Pearson correlation test between real personality scores and predicted one, all five personalities in male manifest significant predictions: E (Pearson’s correlation ρ=0.163, *P value*=0.001), A (Pearson’s correlation ρ=0.1135, *P value*=0.0223), C (Pearson’s correlation ρ=0.206297, *P value*=2.8), N(Pearson’s correlation ρ=0.1475, *P value*=0.0029) and O ((Pearson’s correlation ρ=0.1427, *P value*=0.004) respectively; meanwhile the personality E (Pearson’s correlation ρ=0.1132, *P value*=0.019) in females were significantly better than random (Figure 3). In general, PCA revealed similar association patterns compared to that of CPLSC: males exhibited tentative or significant signals in most personal traits; conscientiousness in males can be best predicted; and females showed substantial correlation only for extraversion. Differences also exist, that associations of E, N and O are strongly significant in PCA but are weak or missing in CPLSC analysis.

**Figure 3.**
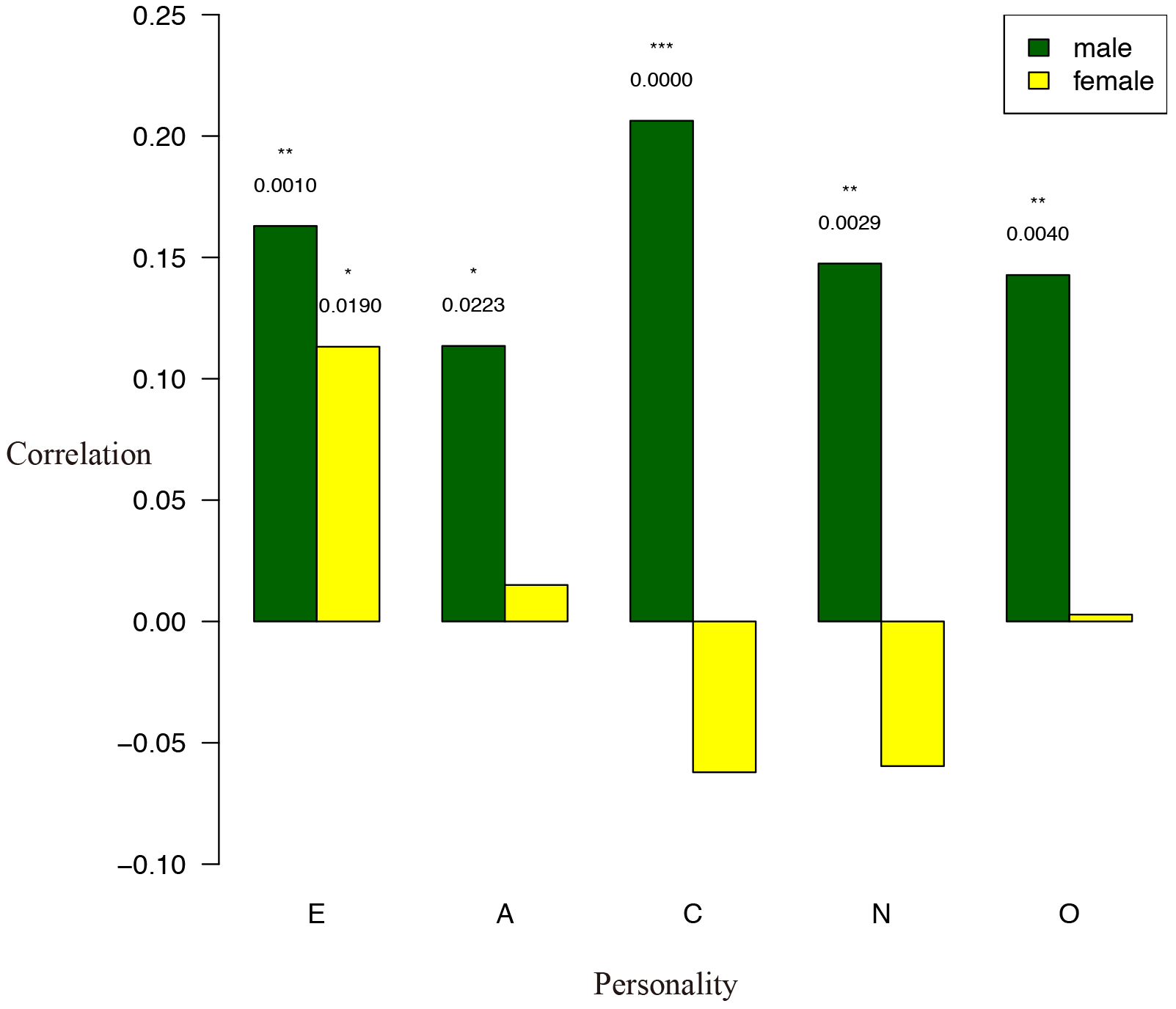
**Pearson correlations between surveyed personality scores and the predicted ones under LOO procedure.** A linear model is used to predict each personality score based on 20PC facial data. P-values and significances of correlation tests are labeled above the bar as numbers and asterisks respectively. *:p<=0.05; **:p<=0.01;***:p<=0.001.

Next, we try to find out the specific facial features that correlated with personalities. We calculated the Pearson correlation in the all-200 combinations of 20 PCs, 5 personalities and two genders. In order to estimate the false discovery rate, we randomly shuffled the sample labels and re-calculated the Pearson correlation 10000 times (see method). The cutoff of correlation p-value thus defined was 0.0033 for the lowest FDR. In total, six PCs were found significantly associated with personality, including in male PC3 and PC16 correlated with extraversion, PC5 and PC15 correlated with agreeableness and PC20 correlated with conscientiousness; and in female, PC4 was correlated with higher extraversion (Figure 4). Moreover, only average 0.676 false-positives were found under permutation (FDR=0.11). It is worth noting that for the shared association signals, CPLSC and PC exhibited similar facial variations. In males, the CPLSC model for agreeableness revealed very similar pattern as PC5, which also showed strongest association to A score (Figure 2 and Figure 4). For conscientiousness in males, the CPLSCand PC20 faces with higher C scores similarly showed tensed jaw muscles (Figure 2 and Figure 4). In females, the PC4 that correlates strongly with E scores showed high similarity to CPLSC model of extraversion (Figure 2 and Figure 4).

**Figure 4.**
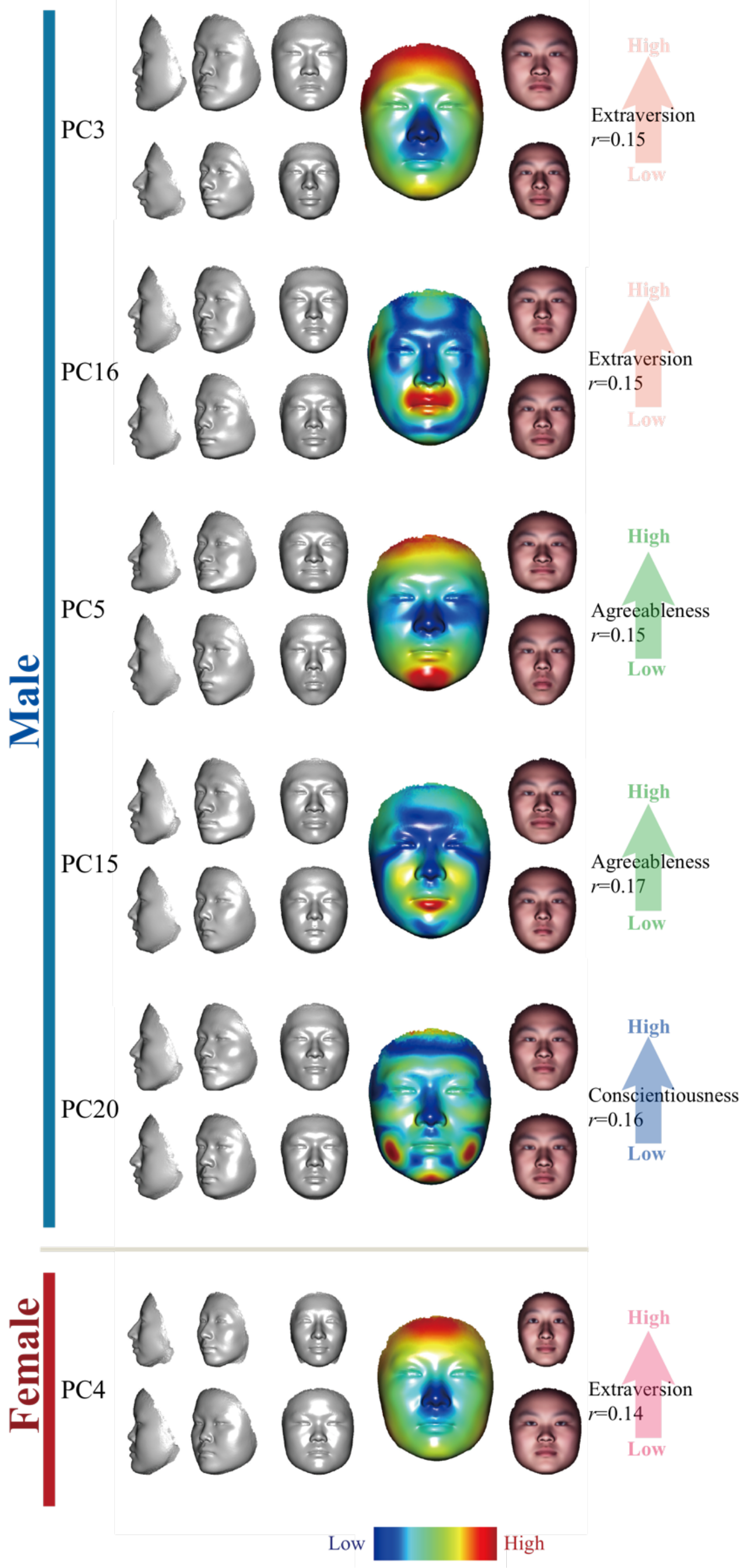
**Extracted personality-related feature which has significant correlation between predicted score and true score by PCA**. From top to bottom are Extraversion-related PC3 and PC16 in male; Agreeableness-related PC5 and PC15 in male. Conscientiousness-related PC 20 in male. Extraversion-related PC14 in female.F each panel,the upper faces are created by adding five standard deviation of projected samples to the mean face, the lower faces are created by subtracting standard deviation of projected samples to the mean face. From left to right are faces rotated by 90°,45° and 0°. The bigger face next is the mean face, on which the heat colors represent the norm value of PCA at each vertex. At the right of the mean face, there are two faces the same as the faces at the left of mean face, but with texture.

## Discussion

Reading personality from face is a fascinating issue. Previous studies based on static 2D images provided evidences that there exist facial traces allowing observers to judge the personality at certain accuracy, but concrete signatures of morphological changes were not obtained. This is mainly due to the difficulty of extracting morphological features from 2D images. With the development of 3D imaging technology and related analysis methods, it is now possible to identify personality related facial features through high density 3D facial image data. In this study, we developed an integrative analysis pipeline: CPLSC, to study the association between 3D faces and personality factors, and extracted effective personality-related features from the image data. To our knowledge, this study provided the first quantitative morphological signatures associated to personality factors. This and further studies of such would provide computational basis, on which stand-alone face recognition platforms may be developed to“read”personality and other psychometric traits from 3D facial images.

We used two methods, namely the CPLSC and PCA to carry out the analysis. In general, there is a clear overlap in the signals identified by both methods. Agreeableness and Conscientiousness in male and Extraversion in female showed statistical significanc and resembled facial features in both CPLSC and PCA analyses. The CPLSC method is based on PLS. PLS is a latent variable approach that is supervised to find the maximal fundamental relationships between the independent variables (e.g. the BF factors) and responses (e.g. the face shape). CPLSC summarizes the face-personality relationships identified by major PLS components, and likely represents the exact facial changes induced by personalities. In comparison, we used a composite PCA model to test the face/personality association, and individual PC components to identify the personality associated facial features. The composite PCA method seems to have a better test power than CPLSC, as all five BF factors were tested significant in male (Fig. 4). For identification of the personality associated features, subtle facial changes due to personality may appear in minor PC components. We dealt with this problem by examining up to 20 PCs, some of which compose only minor fractions of the total variance. The risk of getting more false positive signals in minor PCs was avoidby introducing stringent reshuffling and FDR procedures. On the other hand, as PCA decomposition is unsupervised, the facial variation pertaining to each PC mode may not be exclusively accounted for by personality, but may be partially associated with a BF factor by chance. This could explain why some PC faces of the same BF factor seem to change in opposite directions: in males PC3 and PC16 were both significantly correlated with extraversion/introversion. However, for PC3 the face of higher extraversion scores is wider and more round, whereas for PC16 the extroversive face is associated with a narrower jaw (Fig. 4). The facial signals of extraversion (PC4) in females also seem to reverse the PC3 pattern in males: the thinner face of female PC4 with a pointier nose seems to indicate higher extraversion than the broader face (Fig. 4). These evidences suggest that characteristic features given by PCA may not be very reliable.

Overall, our results suggest that in Han Chinese, male’s personality is more apparent on face than female, supported by both CPLSC and PCA methods. This trend is different from previous studies in European populations [10, 25], suggesting a role of social cultural difference in the exhibition of personality. In our study, Agreeableness, Conscientiousness and Extraversion seem to be more recognizable than other personality traits, as is generally consistent with many previous studies [10, 23, 25, 29]. It would take further detailed studies, especially with much bigger sample sizes, to answer the question whether and why personality traits express on face at different intensities. After all, given the solid evidences of face/personality associations identified in this study, on apure quantitative basis without using human raters, it can be argued that all the BF factors and any other psychological traits may leave some morphological cues on face, which can be“read”by automatic image processing software. To achieve this, bigger sample size is needed; methods should be designed to focus on facial regions that are sensitive to the targeted personality traits; and temporal dimension may be added to capture subtle facial movements related to personality in 4D data.

## Method

### Sample

405 male and 429 female Han Chinese volunteers from Fudan University, Shanghai, took part in 3D image collection and answered the questionnaire to assess their BF personality scores. All the participants are Chinese residents, with males’ age ranging between 16-35 years old (mean: 21.66, sd: 3.69) and female’age ranging between 15-42 years old (mean: 21.65, sd: 3.76).

### 3D image Acquisition and Processing

All the 3D facial images of the sample were collected by high resolution 3D camera system (3dMD face system, www.3dmd.com/3dMDface), then dense non-rigid registration was applied to align all the 3D images according to anatomical homology [30]. After the alignment,each 3D facial image data contains 32251 3D vertices.

### The Big Five Inventory

The Big Five Inventory[31] was downloaded from (http://www.ocf.berkeley.edu/∼johnlab/bfi.htm) and translated into Chinese. This questionnaire is based on the personality model of five factors, including extraversion (E), agreeableness (A), conscientiousness (C), neuroticism (N) and openness (O). The whole inventory is composed of 44 questions in short and easy-to-understand phrases. Each question is designed to be self-rated in a 1 to 5 scale. The Cornbach’s statistic in psychometrics was used to evaluate the robustness of our survey [26].

### Composite Partial Least Square Components

In our study, we proposed the CPLSC framework to integrate the features obtained from individual partial least square components (PLSC)[32]. Briefly, PLS is a kind of dimension reduction method when dealing with the problem that the number of factors is much larger than the sample size, or that the factors are highly collinear. The basic assumption for PLS is that the data observed is produced by a system driven by a few of latent variables[33]. We used the R package“pls” developed by Bjorn-Helge Mevikand Ron Wehrens to carry out the PLS analysis in this study[34, 35].

The CPLSC method begins with determination of the effective number of PLSCs. We introduce coefficient of determination R^2^ to evaluate the performance of model predictions with increasing number of PLSCs. The definition of R^2^ is as follows: assuming we have a set of data *u_i_*, *i*=1,2,3…*n* and corresponding prediction *f_i_*, *i*=1,2,…,n. ū is the mean of *u_i_*, *i*=1,2,3…n. then the total sum of square are:

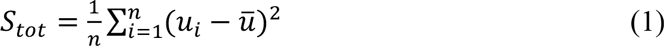
the residual sum of square is: 
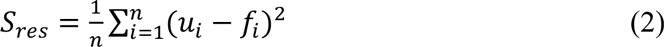
so the definition of R^2^ is: 
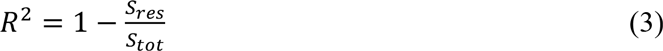

The prediction *f* was obtained by the leave-one-out(LOO) procedure.In brief, given *N* individuals in a sample, one individual is out, anda prediction model is constructed based on the rest *N*‐1 individuals using PLS regression. This model can be used to predict the personality*f_i_* of the left-out individual based on its face data. The LOO procedure was repeated for every individual. The first *m* (*m*=1,2,3…20) PLSCs were used to construct a model, and the optimized m with largest R^2^ value was used as the effective number of PLSCs (Fig. 1).

To formally check the statistical significance of correlations between face and personality, we conduct a permutation procedure. First, for the observed data, we construct a PLS model using the effective *m* PLSCs, and carry out a LOO step as described above to obtain the predicted personality *f_i_* for each individual *i*. As each individual has its true score *u_i_*, the correlation ρcan be calculated between ***f*** and ***u***. Second, we randomly permute the personality scores among different individuals. Based on the permuted sample sets, we can repeat the above pipeline to determine the correlation ρ^*^ under the null hypothesis of no association. The permutation procedure is repeated1000 to give rise to a null distribution Π of ρ^*^. A empirical P-value can be calculated for p in terms of its ranking position in Π.

Given the effective number of PLSCs in the optimal model, we can establish a single linear model CPLSC that combines all the effective PLSCs and describe how a face changes along the BF scores. Given the effective number of PLSCs *m* and each effective *PLSC*_*i*_=(*l_il_*, *l_i2_*,…, *l_ip_*)^T^ *i*=1,2,=,*m*, where *p* is the dimension of all the facial image data, we first normalize the *PLSC*_*i*_ as *NPLSC*_*i*_=*PLSC*_*i*_/‖*PLSC*_*i*_‖, where 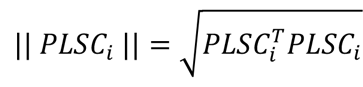 Then we calculate the standard deviation *sd_i_i*=1,2,…, *m*, of the sample for each *NPLSC*_*i*_ *i*=1,2,…, *m*. Finally, the CPLSC model is given as follows:

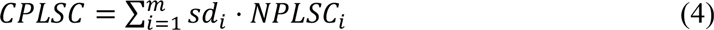

### PCA analysis

PCA is carried out by princomp in MATLAB. A similar LOO procedure is used to evaluate the statistical significance of association based on PCA. Briefly, we used all but one sample to train the model, and use this model to predict the excluded sample. The training-testing procedures were done on every individual. The predicted and real data is then compared.

### R package“3DFace”

To implement the visualization and animation, based on“rgl” package which is 3D graphic utility in R, we developed R package“3DFace”. “3DFace”provides several functions to read the 3D image data, plots the 3D face in different styles and with different gradient color, and generate fantastic 3D animation for the facial feature we extracted. The source code can be download through: https://github.com/fuopen/3dface.

## Reference

1. Peng,S., Tan,J., Hu,S., Zhou,H., Guo,J., Jin,L., & Tang,K. (2013). Detecting genetic association of common human facial morphological variation using high density 3D image registration. PLoS Comput Biol, 9(12), e1003375.

2. Guo,J., Tan,J., Yang,Y., Zhou,H., Hu,S., Hashan,A.,Jin,L.,& Tang,K. (2014). Variation and signatures of selection on the human face. Journal of human evolution, 75, 143–152.

3. Belhumeur,P.N., Hespanha,J. P., & Kriegman,D. J. (1997). Eigenfaces vs. fisherfaces: Recognition using class specific linear projection. Pattern Analysis and Machine Intelligence, IEEE Transactions on, 19(7), 711–720.

4. Hammond,P., Hutton,T. J., Allanson,J. E., Buxton,B., Campbell,L. E., Clayton-Smith,J.,Hespanha,J. P.& Patton,M. (2005). Discriminating power of localized three-dimensional facial morphology. The American Journal of Human Genetics, 77(6), 999–1010.

5. Kramer,R. S., & Ward,R. (2010). Internal facial features are signals of personality and health. The Quarterly Journal of Experimental Psychology, 63(11), 2273–2287.

6. Chen,W., Qian,W., Wu,G., Chen,W., Xian,B., Chen,X.,Ward,R.& Han,J. D. J. (2015). Three-dimensional human facial morphologies as robust aging markers. Cell research, 25(5), 574–587.

7. Little,A. C., Jones,B. C., & DeBruine,L. M. (2011). Facial attractiveness: evolutionary based research. Philosophical Transactions of the Royal Society of London B: Biological Sciences, 366(1571), 1638–1659.

8. Walster,E., Aronson,V., Abrahams,D., & Rottman,L. (1966). Importance of physical attractiveness in dating behavior. Journal of personality and social psychology, 4 (5), 508.

9. Buck,R., Miller,R. E., & Caul,W. F. (1974). Sex, personality, and physiological variables in the communication of affect via facial expression. Journal of personality and social psychology, 30(4), 587.

10. Penton-Voak,I. S., Pound,N., Little,A. C., & Perrett,D. I. (2006). Personality judgments from natural and composite facial images: More evidence for a “kernel of truth” in social perception. Social Cognition, 24 (5), 607–640.

11. Hassin,R., & Trope,Y. (2000). Facing faces: studies on the cognitive aspects of physiognomy. Journal of personality and social psychology, 78 (5), 837.

12. NormanW.T. (1963). Toward an adequate taxonomy of personality attributes: Replicated factor structure in peer nomination personality ratings. The Journal of Abnormal and Social Psychology, 66 (6), 574.

13. Digman,J. M. (1990). Personality structure: Emergence of the five-factor model. Annual review of psychology, 41 (1), 417–440.

14. Goldberg,L. R. (1990). An alternative" description of personality": the big-five factor structure. Journal of personality and social psychology, 59 (6), 1216.

15. De Raad,B. (2000). The Big Five Personality Factors: The psycholexical approach to personality. Hogrefe & Huber Publishers.

16. McCrae,R. R., & Costa,P. T. (2003). Personality in adulthood: A five-factor theory perspective. Guilford Press.

17. Bouchard,T. J., & McGue,M. (2003). Genetic and environmental influences on human psychological differences. Journal of neurobiology, 54 (1), 4–45.

18. Schmitt,D. P., Realo,A., Voracek,M., & Allik,J. (2008). Why can’t a man be more like a woman? Sex differences in Big Five personality traits across 55 cultures. Journal of personality and social psychology, 94 (1), 168.

19. McCrae,R. R., & Allik,I. U. (2002). The five-factor model of personality across cultures. Springer Science & Business Media.

20. Benet-Martinez,V., & John,O. P. (1998). Los Cinco Grandes across cultures and ethnic groups: Multitrait-multimethod analyses of the Big Five in Spanish and English. Journal of personality and social psychology, 75 (3), 729.

21. Weiss,A., King,J. E., & Figueredo,A. J. (2000). The heritability of personality factors in chimpanzees (Pan troglodytes). Behavior genetics, 30 (3), 213–221.

22. Passini,F. T., & Norman,W. T. (1966). A universal conception of personality structure?. Journal of personality and social psychology, 4 (1), 44.

23. Albright,L., Kenny,D. A., & Malloy,T. E. (1988). Consensus in personality judgments at zero acquaintance. Journal of personality and social psychology, 55 (3), 387.

24. Shevlin,M., Walker,S., Davies,M. N., Banyard,P., & Lewis,C. A. (2003). Can you judge a book by its cover? Evidence of self-stranger agreement on personality at zero acquaintance. Personality and Individual Differences, 35(6), 1373–1383.

25. Little,A. C., & Perrett,D. I. (2007). Using composite images to assess accuracy in personality attribution to faces. British Journal of Psychology, 98(1), 111–126.

26. Cronbach,L. J. (1951). Coefficient alpha and the internal structure of tests. psychometrika, 16 (3), 297–334.

27. Schmitt,D. P., Allik,J., McCrae,R. R., & Benet-Martínez,V. (2007). The geographic distribution of Big Five personality traits patterns and profiles of human self-description across 56 nations. Journal of cross-cultural psychology, 38 (2), 173–212.

28. land,D. M., & Thomas,E. V. (1988). Partial least-squares methods for spectral analyses. 1. Relation to other quantitative calibration methods and the extraction of qualitative information. Analytical Chemistry, 60 (11), 1193–1202.

29. Watson,D. (1989). Strangers’ ratings of the five robust personality factors: Evidence of a surprising convergence with self-report. Journal of Personality and Social Psychology, 57 (1), 120.

30. Guo,J., Mei,X., & Tang,K. (2013). Automatic landmark annotation and dense correspondence registration for 3D human facial images. BMC bioinformatics, 14 (1), 1.

31. John,O. P., Naumann,L. P., & Soto,C. J. (2008). Paradigm shift to the integrative big five trait taxonomy. Handbook of personality: Theory and research, 3, 114–158.

32. Wold,H. (1966). Estimation of principal components and related models by iterative least squares. In P.R. Krishnaiaah (Ed.). Multivariate Analysis. (pp.391–420) New York: Academic Press.

33. Rosipal,R., & Kramer,N. (2006). Overview and recent advances in partial least squares. In Subspace, latent structure and feature selection (pp. 34–51). Springer Berlin Heidelberg.

34. Mevik,B. H., & Wehrens,R. (2007). The pls package: principal component and partial least squares regression in R. Journal of Statistical Software, 18 (2), 1–24.

35. evik,B. H., Wehrens,R., & Liland, K. H. (2011). pls: Partial least squares and principal component regression. R package version,2,3.

